# Self-destructive altruism in a synthetic developmental program enables complex feedstock utilization

**DOI:** 10.1101/086900

**Authors:** Robert Egbert, Leandra Brettner, David Zong, Eric Klavins

## Abstract

Cooperation through division of labor underpins biological complexity for organisms and communities. In microbes, stochastic differentiation coupled to programmed cell death drives diverse altruistic behaviors that promote cooperation. Utilizing cell death for developmental multicellular programs requires control over differentiation rate to balance cell proliferation against the utility of sacrifice. However, these behaviors are often controlled by complex regulatory networks and have yet to be demonstrated from first principles. Here we engineered a synthetic developmental gene network that couples stochastic differentiation with programmed cell death to implement a two-member division of labor. Progenitor consumer cells were engineered to grow on cellobiose and differentiate at a controlled rate into self-destructive altruists that release an otherwise sequestered cellulase payload through autolysis. This circuit produces a developmental *Escherichia coli* consortium that utilizes cellulose for growth. We used an experimentally parameterized model of task switching, payload delivery and nutrient release to set key parameters to achieve overall population growth, liberating 14-23% of the available carbon. An inevitable consequence of engineering self-destructive altruism is the emergence of cheaters that undermine cooperation. We observed cheater phenotypes for consumers and altruists, identified mutational hotspots and developed a predictive model of circuit longeivity. This work introduces the altruistic developmental program as a tool for synthetic biology, demonstrates the utility of population dynamics models to engineer multicellular behaviors and provides a testbed for probing the evolutionary biology of self-destructive altruism.

## Introduction

Compartmentalization of function across differentiated cell types was essential to the emergence of complexity in biological systems. Organogenesis in plants and animals^1^, schizogamy in polychaete worms^2^ and germ-soma differentiation in *Volvox* algae^3^ are clear examples of these divisions of labor. Many microbial developmental programs utilize stochastic differentiation and programmed cell death as vital components of population fitness^4^. Selection for programmed cell death has been proposed to drive complex behaviors that delay commitment to costly cell fate decisions^5^, enable adaptation to environmental fluctuations^6^, eliminate competitor species^7^, reinforce biofilm structure^8^ and promote colonization of hostile environments^9^. These behaviors represent divisions of labor between subpopulations of progenitor cells that propagate the species and sacrificial cells that provide a public good, analogous to germ and somatic cell lines in multicellular organisms. Many aspects of the emergence of multi-cellular cooperation and the genetic circuits that control its complexity remain unclear. Limited understanding of the network architectures and stimuli that control developmental gene networks constrains efforts to repurpose them for engineered cell behaviors.

Current engineering paradigms of DNA-encoded cellular logic and feedback control circuits fail to encompass the full suite of behaviors necessary to advance the fields of bioprocessing, bioremediation and cell-based therapeutics. Synthetic microbial consortia have been demonstrated to improve bioprocessing efficiency^10^ or to explore other complex behaviors^11^. A major challenge to engineering microbial consortia is the control of community distributions for complex traits. While syntrophic interactions in defined communities may address some of these needs, they may not be sustainable in environments with fluctuating nutrient or microbial constituents. Further, efficient delivery of protein or small molecule payloads to the environment is constrained by the cell membrane, often requiring the expression of payload-specific pumps or secretion signals. Autolysis triggered by chemical^12,13^ or autoinducer^14^ signals allows release of protein payloads, but prevents applications that may require continuous delivery. Synthetic developmental programs could address these challenges, enabling approaches to create and regenerate microbial communities seeded by individual cells that cooperatively perform complex tasks.

## A synthetic altruistic developmental program

Here we present a synthetic developmental program that implements a germ-soma division of labor to cooperatively digest cellulose. The program links a synthetic differentiation controller with autolysis-mediated enzymatic payload delivery, balancing the rates of stochastic differentiation and programmed cell death to drive overall population growth. We constructed a genetic circuit to create cellobiose consumer cells that produce a sub-population of self-destructive altruists at a controlled rate to enable utilization of cellulose as a sole carbon source through extracellular release of cellulase payloads (Figure 1a).

We refer to the system as SDAc, short for self-59 destructive altruism with a cellulase payload. We implemented SDAc in *Escherichia coli* by engineering a native operon to efficiently utilize cellobiose, introducing a genetic toggle switch tuned to function as a differentiation controller, and constructing a cellulase-lysis payload module to execute the altruist behavior (Figure 1b). We used dynamical systems analysis modeling to identify parameter values critical to achieving overall growth and demonstrated control over the circuit behavior by fine-tuning each parameter.

Using multiplexed mutagenesis and selection, we isolated a strain with a growth rate in cellobiose that is 63% its growth rate in glucose. Though *E. coli* does not natively digest cellobiose, we modified the *chb* operon in a recombinogenic MG1655 derivative^15^ by replacing the native chitobiose-regulated promoter with a strong constitutive promoter^16^. We further improved growth on cellobiose by subjecting the constitutive *chb* expression variant to multiple cycles of multiplexed recombineering targeting the *chb* genes and selected for cellobiose utilization in minimal cellobiose media (Supplementary Figure 1). We identified the variant with the highest growth rate, DL069, as a *chbR* deletion mutant.

To control differentiation rate, we constructed and sampled from a library of mutual inhibition toggle switch variants that exhibit regular stochastic state transitions. While genetic toggle switches are often designed to function as bistable memory devices^17^, a quasi-steady state can be achieved by properly balancing expression levels of the repressor proteins^18^. Simple sequence repeats embedded in the ribosome binding site (rbSSR) allow predictable modulation of translation initiation rate to tune the balance between transcriptional repressors^19^.

We engineered altruist payload delivery by constructing a cellulase and lysis gene cassette. The operon was designed to maximize production of the cellulase payload with an efficient ribosome binding site and a poly-`(AT)` rbSSR to fine-tune expression of the lysis gene. In order to minimize the altruist subpopulation it is desirable for maximal post-differentiation accumulation of the payload to precede autolysis. Colicin gene networks share this trait, using stochastic gene expression of colicin and lysis genes within subpopulations to kill ecological competitors^7,20^. We found that coupling the lysis gene from colicin E3 to the differentiation controller enabled stochastic state transitions and delayed lysis at the microcolony level, evidenced by accumulation of a GFP payload followed by autolysis (Figure 1c and Supplementary Movie).

**Figure 1:**
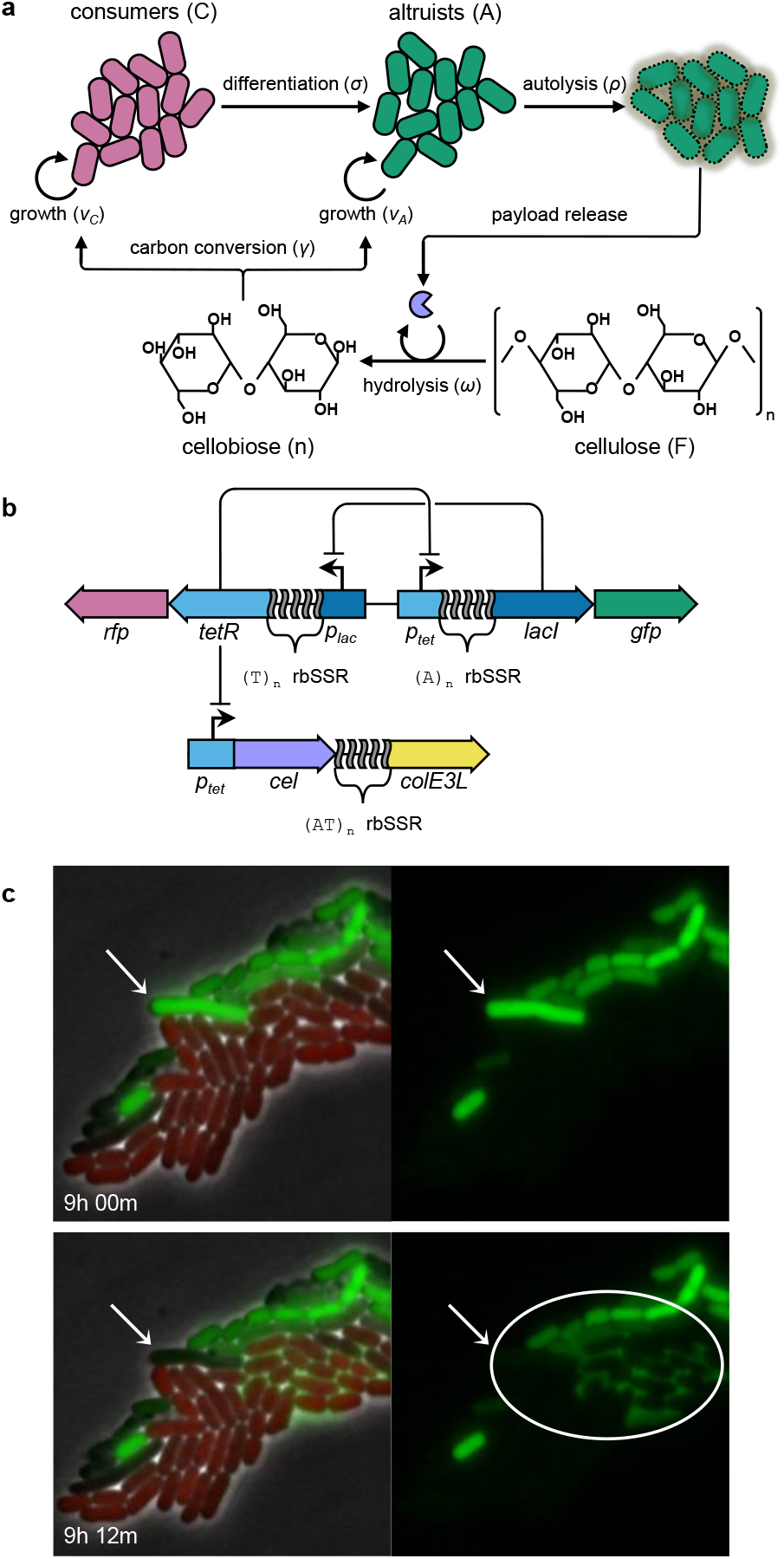
A synthetic developmental program for cooperative cellulose digestion. (a) Cellobiose consumers stochastically transition to self-destructive altruists. Altruists, in turn, produce and release cellulase payloads via autolysis to support the consumer population. (b) The genetic implementation of the SDAc developmental program includes a differentiation control plasmid (above) and a payload delivery plasmid (below). Cell states are mediated by a mutual inhibition toggle switch using transcriptional repressors LacI and TetR. TetR-dominant cells express RFP as consumers; LacI-dominant cells co-express cellulase and colE3 lysis (*colE3L*) proteins with GFP as altruists. Differentiation and lysis rates are fine-tuned with rbSSR sequences for *tetR* and *lacI* (differentiation) and *colE3L* (lysis). (c) Demonstration of differentiation and autolysis within a microcolony seeded by a single cell. A large altruist cell (arrow) accumulates its GFP payload (upper panel) until it undergoes autolysis (lower panel), enabling payload diffusion to surrounding cells (circle). See Supplementary Movie.

## SDAc parameter estimates through modular system decomposition

Analysis of a population dynamics model of SDAc behavior suggested optimal parameter regimes for cellulose utilization and guided implementation of the developmental circuit. Though we observed the requisite behaviors of differentiation, payload accumulation and autolytic payload release at the microcolony level, it was not clear what combination of expression levels for circuit components would enable cooperative growth on cellulose. To reason about the functional parameter space for SDAc behavior we developed a population dynamics model using a system of ordinary differential equations that maps system parameters to experimentally tunable features of the genetic circuit. The model species are consumers, altruists, cellulose feedstock, and cellulose-derived nutrients. These species and the associated kinetic parameters are described in Box 1.

#### Box 1. Systems of equations for modeling synthetic self-destructive altruism.

I. 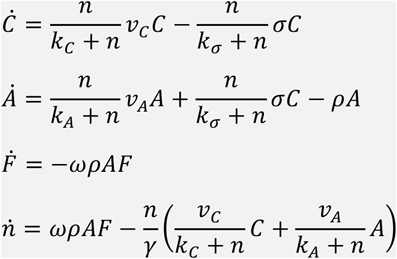
II. 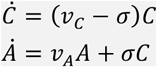
III. 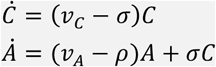
IV. 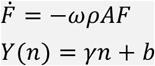
V. 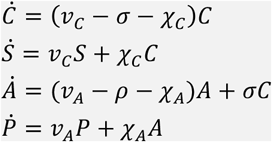

**Module I**. We constructed a population scale model composed of first order ordinary differential equations. The model contains four relevant species: consumers (*C*), altruists (*A*), cellulose (*F*, feedstock), and digestible nutrients (*n*) with corresponding units of colony forming units per mL for cells and grams per mL for molecules. Consumer and altruist cells grow in the presence of nutrients at rates *v_C_* and *v_A_*, respectively. Individual consumer cells differentiate to altruists at rate *σ*, and altruists lyse at rate *ρ*. Altruist payloads degrade feedstock to nutrients at rate *ω*, and nutrients yield biomass according to *γ*. Nutrient-dependent dynamics are controlled by half maximal rate constants for growth (*k_C_, k_C_*) and differentiation (*k*_*σ*_). Autolysis is considered nutrient-independent. Additional model details are included in Supplementary Table 1 and Supplementary Note 1.

#### Experimental Parameter Measurements

The modularity of the synthetic SDAc developmental gene network allows experimental measurement of each parameter by systematic deconstruction of the full system. We constructed simplified sub-models to identify the key circuit parameters and measured the behaviors of defined sub-circuits to estimate the parameters. Experimental details are described in the Materials and Methods and modeling approaches to the parameter estimates are described in detail in the Supplementary Notes.

**Module II**. Differentiation rate (*σ*) A continuous growth model of consumer and pseudo-altruists was used to estimate differentiation rate for SDAc strains missing payload and lysis genes. The temporal population fraction of consumers (RFP producers) and altruists (GFP producers) was measured by flow cytometry for strains pre-induced to the consumer state, washed and grown in M9 minimal cellobiose media (Figure 2a–c). For each strain *σ* estimates were fit to this system of equations using growth rates measured independently. Unbounded growth in the dynamics represents growth for each passage over a finite duration of the periodic dilution.
**Module III**. Autolysis rate (*ρ*) A continuous growth model of consumers and altruists was used to estimate autolysis rate for SDAc strains. Cultures were initialized and measured as in Module II. For each strain *ρ* estimates were fit to this system of equations (Figure 2d–f) using growth rates and differentiation rates measured independently.
**Module IV**. Cellulose hydrolysis (*ω*) A model of feedstock degradation and nutrient release was used to estimate cellulose hydrolysis rates for lysis deficient SDAc strains expressing cellulase payloads. Crude cell lysates were generated from cellulase producing strains. Nutrient release from PASC media inoculated with lysates was measured via supernatant growth of a cellulase and lysis deficient strain. For each strain *ω* estimates were fit to the feedstock equation *F* using nutrient estimates derived by applying the equation for *Y* to growth data.
**Module V**. Cheater dynamics A continuous growth model of consumers, altruists, consumer cheaters that do not differentiate (*S*) and altruist cheaters that do not lyse (*P*) was used to estimate rates of escape from consumer (χ*_C_*) and altruist (χ*_A_*) states. Models that account for single or dual cheater subpopulations were used as fits to the population fraction data and growth rate data obtained for Module II (Figure 4d).

The core tunable parameters for SDAc cellulose utilization are the growth rate on cellobiose, differentiation rate, lysis rate and cellulase activity. Cells must utilize the hydrolysis products of the cellulase enzymes, including cellobiose. Insufficient differentiation would limit growth via low cellulase release, while excessive differentiation would incur unnecessary fitness defects for consumers or, at extreme rates, to population collapse. Low lysis rates would limit feedstock degradation through sequestration of intracellular cellulase and rapid lysis would reduce the per-altruist payload burst size. High cellulase activity improves growth titer by reducing the population fraction of altruists required to deconstruct the feedstock. Ultimately, SDAc performance is constrained by the tunability of the circuit components and many parameter sets predict no growth on cellulose (Supplementary Figure 2).

The modularity of a synthetic gene circuit implementation allowed us to decompose the system model and its experimental components to estimate system parameters and predict cellulose utilization for the full circuit. We developed the modules described in Box 1 to measure growth rate, differentiation rate and lysis rate in cellobiose as well as cellulase activity, drawing from a small parts library for each module to sample a range of parameters.

We experimentally tuned differentiation rate over an order of magnitude with a collection of SDAc strains lacking cellulase and autolysis genes. Specifically, we used multiple poly-`(T)` rbSSR variants controlling expression of the consumer-dominant regulator TetR to modulate the differentiation rate from consumer to LacI-dominant altruists (Figure 2a,b). We measured the population fraction of differentiated cells as a function of time using flow cytometry (Supplementary Figure 3) and fit a two-state, continuous growth model to the data for consumers transitioning to altruists at rate *σ* (Box 1, Module II). We found the repeat length to be inversely proportional to differentiation rate, supporting previous results for a switch on a higher copy number plasmid^19^ and resulting in *σ* estimates ranging from 2.7×10^−2^ ± 9.0×10^−3^ *h*^−1^ for (T)_12_ to 2.11×10^−1^ ± 1.5×10^−2^ *h*^−1^ for (T)_18_ (Figure 2c, Supplementary Table 4).

Lysis rates were modulated over a four-fold range using expression variants of the colicin E3 lysis gene. Using the intermediate rate differentiator (T)_16_, we tested a set of poly-`(AT)` rbSSR variants to modulate lysis gene expression (Figure 2d,e). We used the same population fraction assay as for differentiation to fit a consumer growth and differentiation model that includes altruist lysis parameter *ρ* (Box 1, Module III). We measured lysis rates from 7.2×10^−2^ ± 1.8×10^−2^ ℎ^−1^ for (`AT`)_10_ to 1.9×10^−1^ ± 2.5×10^−2^ ℎ^−2^ for (`AT`)_8_ (Figure 2f, Supplementary Table 5). As predicted by the differentiation with autolysis model, the differentiated population fraction for each switch variant with the lysis gene is lower than for the equivalent autolysis-178 deficient strain (Supplementary Figure 5). We found, however, that the lysis rate did not correlate with rbSSR length (Supplementary Figure 6).

To estimate cellulase activity we quantified cellulose degradation from cell lysates of autolysis-deficient SDAc strains producing one or two cellulases, observing hydrolysis rates over a three-fold range. We measured cellulose degradation and digestible nutrient release for three endoglucanases from two glycoside hydrolase (GH) families: CelD04^21^ and BsCel5^22^ from GH5; and CpCel9 from GH9^23^ (Figure 2g,h). We also measured the activity of multi-enzyme cocktails using each GH5 enzyme with CpCel9, combinations with reported synergistic activities^24^. We used Congo Red staining of M9 minimal phosphoric acid swollen cellulose (PASC) media spiked with cell lysate to observe cellulose degradation up to 23% (Supplementary Note 6) in and quantified cell growth on the resulting supernatant to estimate nutrient release of up to 14% of cellobiose equivalents (see Supplementary Figure 8, Supplementary Note 6). We used these cellulase activity measurements to fit a value for *ω* to the feedstock differential equation (Box 1, Module IV). Cellulase activity estimates range from 6.0 × 10^−13^ CFU^−1^ mL for CpCel9 to 1.9 × 10^−12^ CFU^−1^ mL for a BsCel5/CpCel9 cocktail (Figure 2i, Supplementary Table 7).

**Figure 2.**
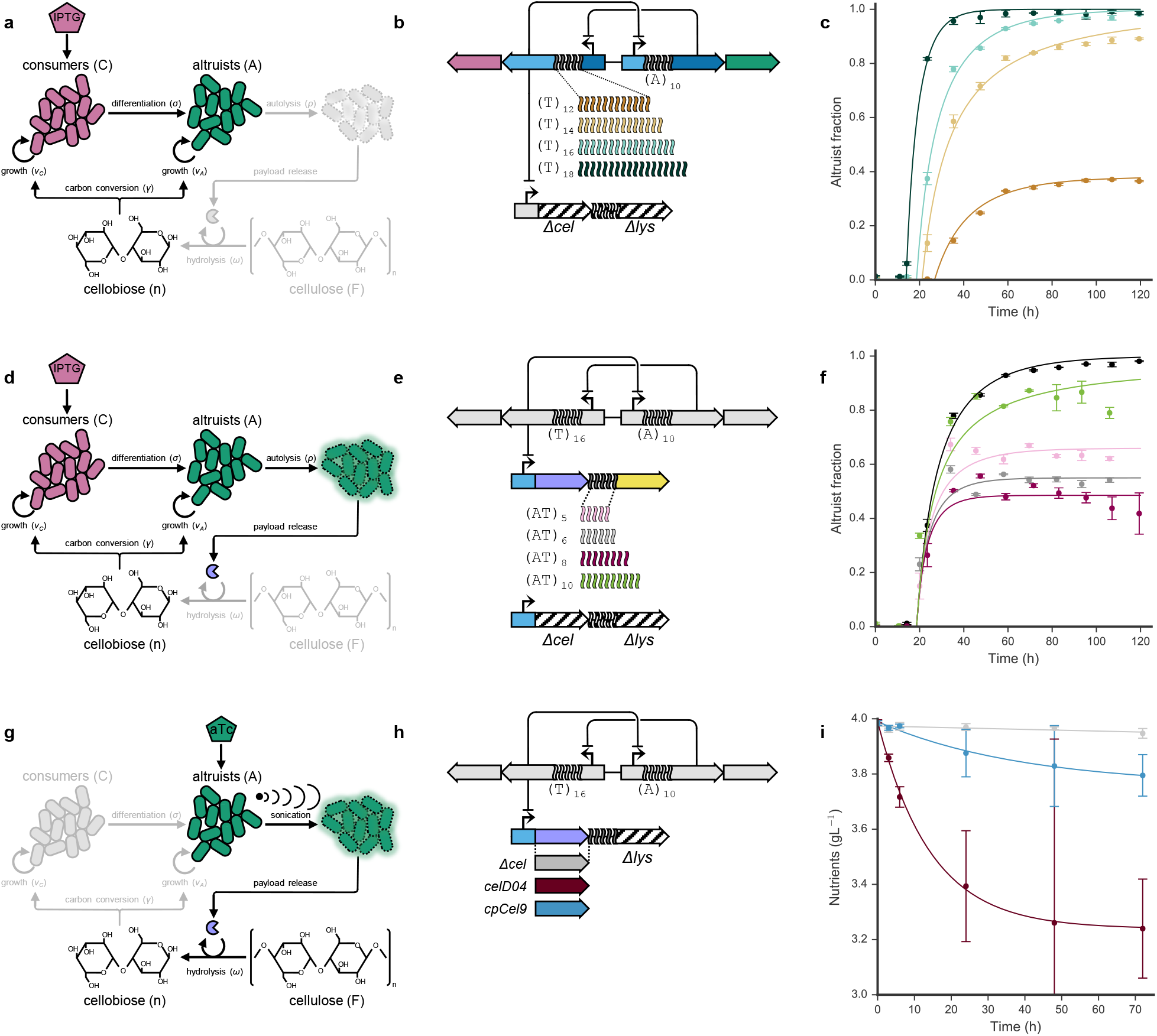
Experimental parameter sweeps for differentiation, autolysis and cellulase activity. (a) For the differentiation assay, cultures were initialized to the consumer state using IPTG and grown continuously in cellobiose to differentiate into cellulase-and lysis-deficient altruists. Cartoon components in grey correspond to unutilized states for the assay. (b) Genetic variants tested for differentiation vary poly-`(T)` rbSSR length to modulate TetR expression. Cellulase and lysis genes were removed from the payload plasmid (Δ*cel*, Δ*lys*). (c) Differentiation data and model fits to estimate *σ* (see Supplementary Note 4). Plot colors correspond to constructs depicted in (b). (d) For the lysis assay, cultures were initialized as in (a) for consumers to differentiate into autolytic altruists. (e) Genetic variants tested for lysis used intermediate rate differentiator (T)_16_, varying `(AT)`-rbSSR repeats that control lysis gene expression or using a control plasmid with no cellulase or lysis genes (Δ*cel*, Δ*lys*). (f) Lysis data and model fits to estimate *ρ* (see Supplementary Note 5). Control from (c) shown in black for comparison. (g) For the cellulase activity assay, lysis-deficient strains were induced to the altruist state using aTc, grown to saturation and sonicated to generate crude cell extracts. (h) Genetic variants tests for cellulase activity by expressing different cellulases or maintaining a control plasmid (Δ*cel*). (i) Cellulase activity data and model fits to estimate *ω* (see Supplementary Note 7).

## Cellulose utilization with full circuit model predictions

To quantify the combined effects of differentiation and autolysis dynamics on feedstock degradation and cell growth we measured cellulase activity from SDAc strains with the full circuit. Cellulose hydrolysis by individual colonies was measured by Congo Red clearing assays from agar plates supplemented with carboxymethylcellulose (Figure 3). We found that the clearing diameter for switch variants increased as a function of differentiation rate (Figure 3a,b). We observed no clearings for a control lacking cellulase. We also tested the effect of rbSSR lysis variants combined with cellulase CelD04 (Figure 3c,d) as well as for individual cellulases (Figure 3g,h) using intermediate rate differentiator (T)_16_. We found the GH5 cellulases generated larger clearings than CpCel9, consistent with the in vitro cellulase activity results.

Fine-tuning the differentiation, lysis and cellulase activity parameters is critical to realizing robust SDAc growth on cellulose as a sole carbon source. To determine fitness on cellulose and validate the full dynamics model (Box 1, module I), we measured viable cell counts in PASC for SDAc variants that span a range of values for each core parameter. We observed the highest population fitness at intermediate differentiation rates (Figure 3c), with high lysis rates (Figure 3f) and with high cellulase activity (Figure 3i), trends that are consistent with the naive model predictions from Supplementary Figure 2. Model fits of growth dynamics using parameter estimates from individual modules match observations for most variants, though the model predicted higher growth for differentiation variant (T)_18_ and failed to capture growth lag dynamics for the BsCel5-CpCel9 cellulase cocktail (Supplementary figure 9).

**Figure 3.**
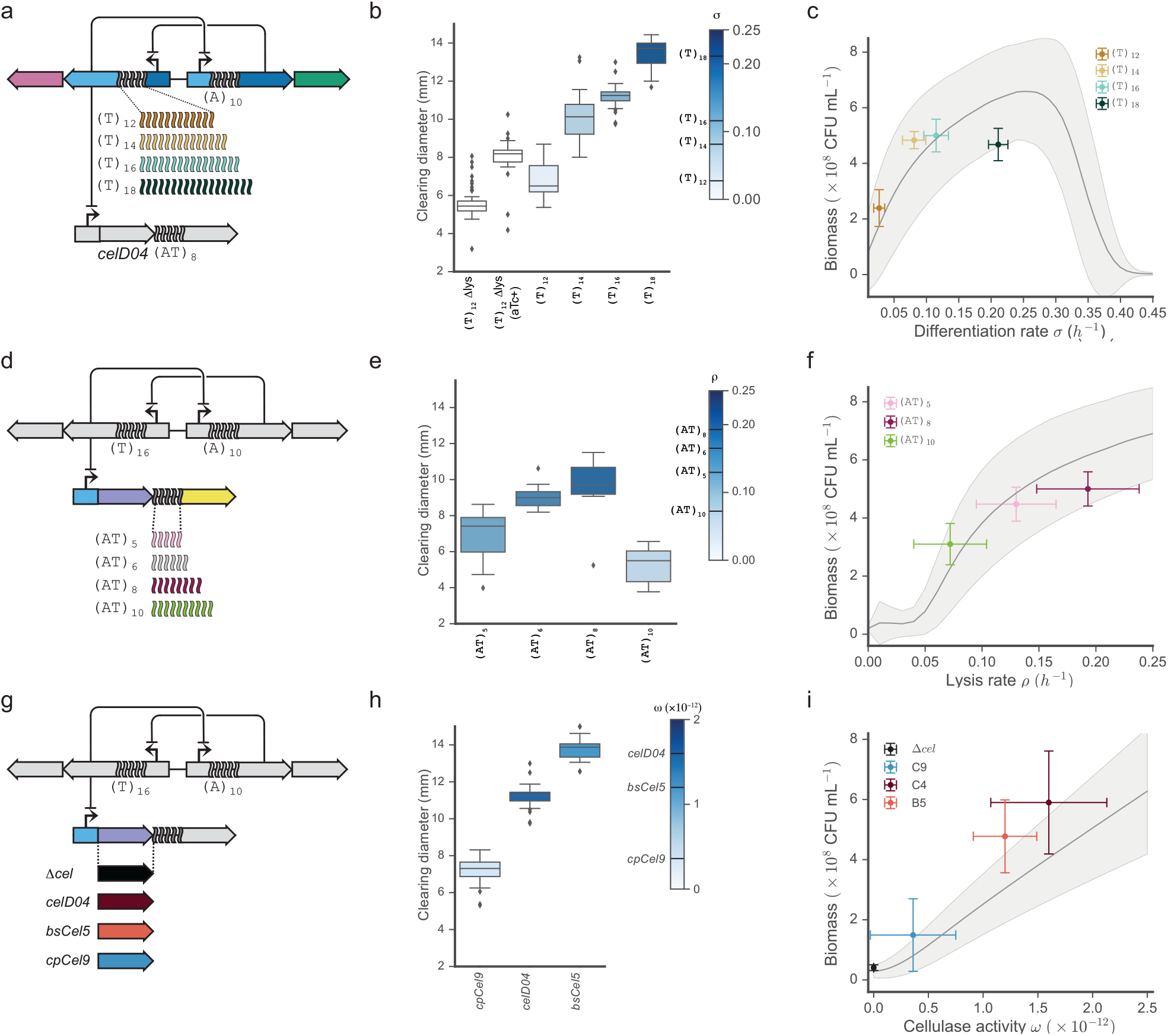
Characterization of cellulose hydrolysis and utilization for growth reveals SDAc model prediction accuracy. (a) Differentiation rate variants expressing cellulase CelD04 with lysis rbSSR (AT)_8_. (b) Clearing size distributions for individual colonies from differentiation variants in (a) (N = 31, 21, 17, 15, 6). (c) Growth titer in M9 minimal 0.4% PASC media after hours for differentiation variants in (a). Error bars on the x-axis represent standard deviation of the parameter estimate and error bars on the y-axis represent standard error for CFU counts from at least four replicates. The shaded region represents one standard deviation of model uncertainty for cell growth. (d) Lysis rate variants for intermediate differentiator (T)_16_ expressing cellulase CelD04. (e) Clearing size distributions for individual colonies from lysis variants in (d) (N = 20, 14, 11, 15). (f) Growth titer as in (c) for some lysis variants in (d). (g) Cellulase variants for intermediate rate differentiator (T)_16_. (h) Clearing size distributions for individual colonies from cellulase variants in (g) (N = 20, 15, 16). (i) Growth titer as in (c) for cellulase variants in (g). Error bars for the x-axis here represent the interquartile range for each *ω* estimate. Whiskers shown for box plots in (b,e,h) extend one interquartile range.

## Excessive differentiation leads to a tragedy of the commons

The full model of SDAc growth dynamics on cellulose predicts system collapse at high differentiation rates (Figure 3a), but does not account for mutational dynamics that could generate non-cooperative cheaters. Indeed, we observed functional instability for hyperdifferentiator switch variant (T)_18_. The instability was manifest in continuous cellobiose culture as a temporally unstable altruist fraction (Supplementary Figure 5). Growth rate dynamics consistent with enrichment for mutants that overcome the population growth rate reductions imposed by differentiation or lysis (see Figure 4e) support this hypothesis. We also observed two mutant colony phenotypes for the same strain after extended growth in PASC media (Figure 4a), further suggesting functional instability at extreme differentiation rates.

Analytical solutions to candidate dynamic models incorporating non-cooperative mutants suggests that two cheater subpopulations – one deficient in differentiation and the other deficient in lysis – are required to realize the observed dynamics. To estimate SDAc mutational rates we developed a model that introduces new species for switch-deficient cheaters (S) and for lysis-deficient pseudo-altruist cheaters (P) and their respective escape rates, χ*_C_* and χ_*A*_ (Box 1, module V). Four candidate models were investigated to account for the cheater dynamics: χ_*C*_ = χ*_A_* = 0 (lysis cheaters) χ*_A_* = 0, χ*_C_* > 0 (differentiation cheaters), χ*_C_* = 0, χ*_A_* > 0 (lysis cheaters) or χ*_C_* = 0, χ*_A_* > 0 (dual cheaters) (Figure 4b). We derived analytical solutions for each model, finding that the only model that supports the observed dynamics includes both cheater types (Supplementary Note 11). Cheaters may emerge from discrete mutational events during growth or be part of the inoculum, rising in population fraction once a large fraction of cells differentiate and lyse. Our analytical solutions do not distinguish between either initial condition.

Fits of each mutagenesis model to measured differentiation and growth dynamics validate the dual cheater model and give mutation rate estimates for each cheater type. We fit each cheater model to the differentiation and lysis data in cellobiose for hyperdifferentiator (T)_18_. Fits for differentiation and growth dynamics are shown in Supplementary Figure and rate estimates are shown in Supplementary Table 9. The lysis cheater model provided no improvement to the null case and is not shown. The dual cheater model fit estimates the emergence of differentiation cheaters at a rate of 1.8×10^−6^ ± 6.3×10^−7^ *h*^−1^ and the emergence of altruist cheaters at a rate of 1.9×10^−5^ ± 8.2×10^−6^ *h*^−1^. Using the mutagenesis parameters fit from hyperdifferentiator (T)_18_, intermediate differentiator (T)_16_ is also predicted to accumulate cheaters within the measurement interval (Figure 4c), which is consistent with the trend of the cellobiose switching data. When applying escape rate estimates to a model of the overall population growth dynamics for differentiation rate variants, we found the model predicted the variable growth rate dynamics observed for variants with high differentiation rates (Figure 4d).

DNA sequencing of cheater isolates confirms the genetic basis for both differentiation and lysis cheaters (Figure 4e). We observed large colonies that were bright red or bright green – putative differentiation and lysis cheaters, respectively – in addition to the wild-type small, mixed-color colony on solid media after extended growth in PASC media. Sequence analysis of the differentiation controller plasmid from red escape colonies isolated from six replicate cultures revealed mutations to two hypermutable loci with predictable effects (Supplementary Table 10): expansion or contraction of the tandem (CTGG)_3_ mutational hotspot observed in four of six replicates should prevent altruist emergence through inactive, truncated Lac repressor^25^; and deletions within the (T)_18_ rbSSR controlling TetR expression (one of six replicates) should abolish differentiation by reducing *σ*. A transposition event of insertion sequence IS2^26^ internal to *lacI* (one of six replicates) should also prevent differentiation. Sequence analysis of the payload delivery plasmid revealed a transposition event of IS1^27^ between the cellulase and lysis genes in one of six sequenced replicates, likely disrupting operon expression (Supplementary Table 11). The majority of the altruist cheater colonies we sequenced revealed no mutations in the payload delivery transcription unit, suggesting lysis evasion via mutations on the genome or elsewhere on the plasmids. Given that the lysis gene is sourced from a colicin plasmid found in natural *E. coli* populations, it is possible the genome encodes high-rate evolutionary paths to lysis immunity.

**Figure 4.**
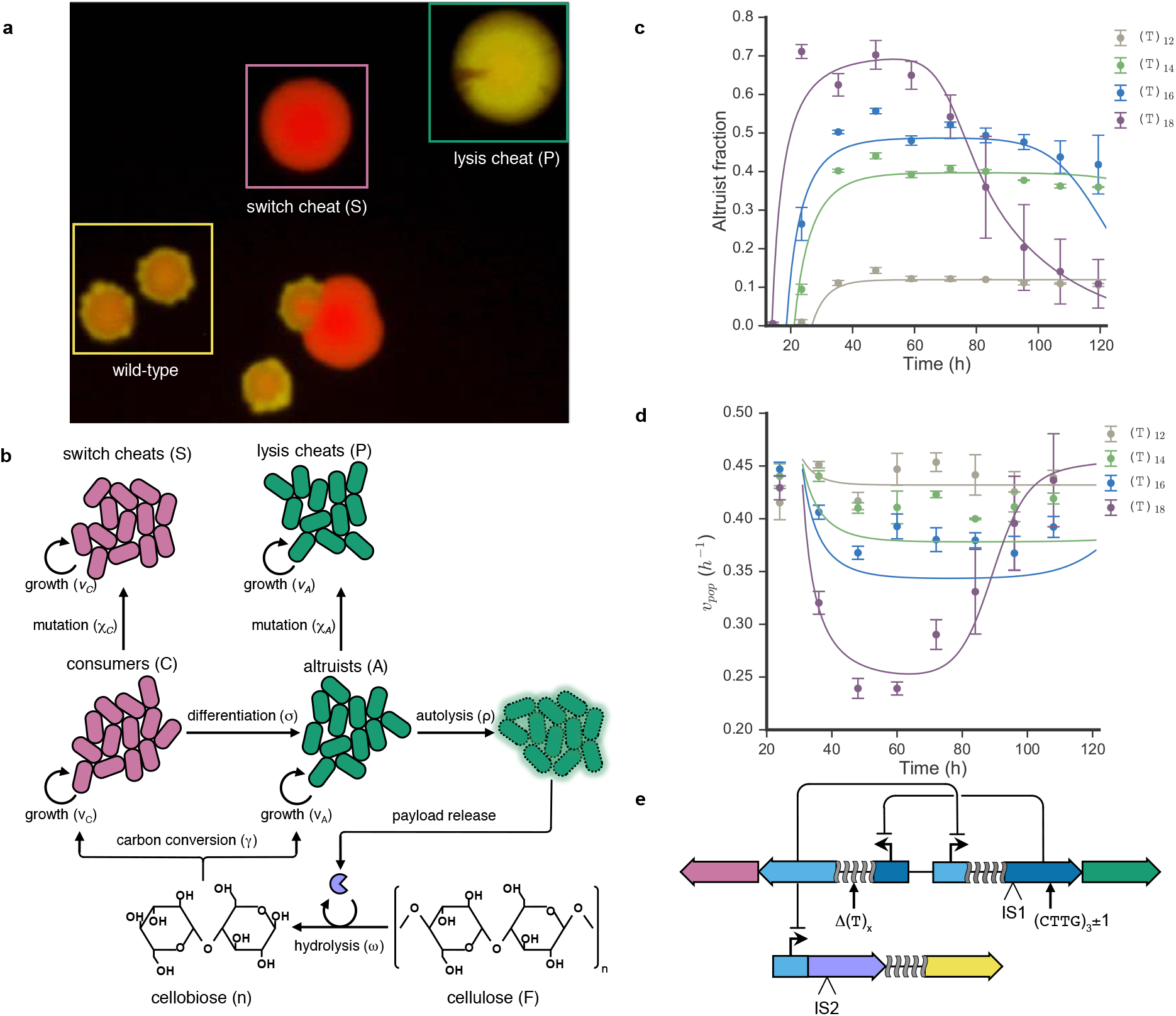
Mechanisms and rates of escape for SDAc cheaters. (a) Fluorescence image of wild-type and cheater colonies isolated from PASC cultures of hyperdifferentiator (T)_18_ that no longer differentiate (S) or no longer lyse (P). (b) Representation of alternate SDAc model that incorporates mutational dynamics by including sink states for switch cheaters (S) and pseudo-altruist cheaters (P) with associated mutagenesis rates (see Box 1, module V). (c) Dual cheater model fit to altruist fraction measurements for differentiation (compare to Supplementary Figure 5a). Escape rate estimates for hyperdifferentiator (T)_18_ are used to fit altruist fractions for the other differentiation rates. (d) Continuous growth model fit for overall population growth rate measurements of differentiation variants (*v_pop_*), using dual cheater rate estimates as in (c). (e) Summary of observed mutations that produce differentiation or lysis cheaters. Differentiation mutants included an expansion or contraction of a native simple sequence repeat element in the *lacI* coding sequence, repeat unit truncations of rbSSR (T)_18_ that controls differentiation rate and insertion sequence disruption of LacI expression. Insertion sequence disruption of the CelD04 gene (green arrows) was observed for altruist cheaters. See Supplementary Tables and for additional details on observed mutations.

## Discussion

We have demonstrated a first-principles approach to construct a developmental gene circuit and have implemented a two-member developmental system to cooperatively utilize the complex feedstock cellulose. In-depth system deconstruction and characterization enables model-guided optimization of growth on cellulose. At extreme differentiation rates, genetic instability drives the emergence of cheaters that fail to differentiate or fail to lyse. This foundation will enable development of more robust and complex developmental divisions of labor to advance sustainable bioprocessing and cell-based therapeutics. Similar systems may also prove to be effective tools to advance the study of the evolution of cooperation.

Due to the observed functional instability for some variants, the SDAc program likely suffers from a tragedy of the commons^28^. In well-mixed cellulose media, emergent cheaters fully benefit from the public good provided by the altruists. Further, due to the costs of switching and lysis, the cheaters can out-compete cooperators and sweep the population. In the absence of altruists, cellulase release ceases, driving population collapse. Previous work has shown that when the environment is spatially organized a communal benefit applies only to nearby, closely related cells who are likely fellow cooperators^29^. Indeed, research suggests the cellulosome evolved to localize the benefits of cellulase expression, as in sucrose utilization in yeast^30^. Thus cheaters are stranded with limited or no access to the shared resource. This phenomenon, attributed to kin selection, could preserve cooperative behavior for many more generations, potentially avoiding the functional instability we observed. Future work could elucidate the role of structured environments in this synthetic system to reduce the impact of cheaters or to evolve more stable cooperator phenotypes.

While we only observed a high fraction of SDAc cheaters from hyperdifferentiation variant (T)_18_, engineering developmental circuits for deployment in bioreactors or other complex environments would require long-term evolutionary stability to minimize cheaters and maintain engineered function. Interestingly, previous studies have shown that *lacI* tandem repeat mutations occur at a rate > 10^−6^ events per generation^25^ and transposon insertion elements jump at rates of 10^−6^−10^−5^ insertions per generation^31^. These rates are consistent with our experimental estimates for mutagenesis, suggesting relatively simple modifications may considerably boost SDAc circuit longeivity. Analysis of the cheater model suggests that a reduction of cheater rates by 100-fold and 1000-fold for intermediate differentiator (T)_16_ would increase circuit stability by 56% and 85%, respectively, boosting the functional period in continuous culture from 2.8 days to 5.1 days. Genetic strategies to boost evolutionary stability include recoding the repeat region of *lacI*, introducing stabilizing degeneracy into rbSSR sequences and porting the system to a low mutation rate strain deficient for insertion elements^32^. Further gains in system performance could be achieved by chromosomal integration of the SDAc network to prevent the fixation of mutant plasmids in the population^33^ or plasmid loss. Finally, incorporating more efficient cellulase cocktails will reduce evolutionary pressure for cheating by decreasing the optimal altruist load.

The division of labor system outlined here is a template for the construction of other developmental programs to perform complex tasks in engineered microbial communities. This work can be extended in many ways. For SDAc, the developmental program could be triggered in response to nutrient depletion when the supply of simple sugars is depleted. Alternative protein and small molecule payloads from a general autolytic delivery system could be designed to mediate microbial interactions, aid in bioprocessing or bioremediation or as a cellular therapeutic. Further, stochastic strategies could be employed with or without self-destructive altruism to seed multicellular developmental programs for distributed metabolic engineering^34^, evolutionary engineering^35^ or to control distributions of multiple cell types in microbial communities^36,37^. Tunable developmental programs could also be applied to better understand the emergence and persistence of well-342 studied developmental programs, substituting complex regulatory networks with tunable differentiation dynamics.

## Acknowledgements

We thank Ben Kerr for helpful discussions regarding evolutionary dynamics of self-destructive altruism and for sharing a sample of the colicin E3 plasmid. We also thank Chris Takahashi for providing strain CT009. This work was funded by National Science Foundation Molecular Programming Project grants 0832773 (R.G.E and E.K.) and 1317653 (L.M.B and E.K.), National Science Foundation Bio/computational Evolution in Action CONsortium (BEACON) grant 0939454 (L.M.B. and E.K.) and the University of Washington Mary Gates Research Scholarship (D.Z.). R.G.E. and E.K. conceived and designed the study. R.G.E., L.M.B. and D.Z. performed the experiments and analyzed the data. L.M.B., R.G.E. and E.K. performed the computational modeling and analyzed simulations. All authors contributed to the manuscript.

